# Lying in a 3T MRI Scanner Induces Neglect-Like Spatial Attention Bias

**DOI:** 10.1101/2021.06.07.447306

**Authors:** Axel Lindner, Daniel Wiesen, Hans-Otto Karnath

**Author notes:** Shared Correspondence (;). All authors contributed equally to the paper.

## Abstract

The static magnetic field of MRI scanners can induce a magneto-hydrodynamic stimulation of the vestibular organ (MVS). In common fMRI settings, this MVS effect leads to a vestibular ocular reflex (VOR). We asked whether – beyond inducing a VOR – putting a healthy subject in a 3T MRI scanner would also alter goal-directed spatial behavior, as is known from other types of vestibular stimulation. We investigated 17 healthy volunteers, all of which exhibited a rightward VOR inside the MRI-scanner as compared to outside-MRI conditions. More importantly, when probing the distribution of overt spatial attention inside the MRI using a visual search task, subjects scanned a region of space that was significantly shifted towards the right. An additional estimate of subjective straight-ahead orientation likewise exhibited an MVC-induced rightward shift. Hence, putting a subject in a 3T MRI-scanner induces a bias of spatial attention, which closely mimics that of stroke patients with spatial neglect.

## Introduction

Recently, Roberts and colleagues (2011) observed that healthy subjects who were exposed to the static magnetic field of 3T and 7T MRI scanners developed a persistent horizontal nystagmus in complete darkness. Since then, this effect was replicated by various groups (e.g. Mian et al 2013, Boegle et al 2016) and was shown to be present for standard fMRI settings at 3T (Boegle et al 2016). Moreover, these studies have further corroborated the initial conclusion by Roberts et al. (2011) that the nystagmus is caused by Lorentz forces that result from the interaction of the static magnetic field of the scanner and ionic currents in the endolymph fluid of the subject’s labyrinth (see Ward et al 2019 for review). This force acts predominantly upon the cupula of the horizontal semicircular canal, inducing a slow horizontal vestibular ocular reflex (VOR) that is accompanied by fast resetting saccades in the opposite direction (Otero-Millan et al 2017). In natural settings, a stimulation of the horizontal canals occurs, for instance, when our head is accelerated to the left or to the right. The resulting VOR then smoothly moves our eyes in the direction opposite to the head rotation. The VOR thus supports – together with the visually induced optokinetic reflex – the stabilization of the retinal image despite head-motion (for review e.g. see Ilg 1997, Angelaki & Cullen 2008).

Roberts and colleagues (2011, p. 1638) pointed out that the fake head-motion signal induced by magneto-hydrodynamic vestibular stimulation (MVS) “carries important ramifications and caveats for functional MRI studies, not only of the vestibular system but of cognition, motor control, and perception in general”. For instance, Boegle and colleagues (2016) observed that MVS does indeed modulate fMRI-signal fluctuations. This modulation affected the default mode network as a function of the strength of the magnetic field and mainly in areas associated with vestibular and ocular motor function. Thus, MVS induces balance shifts in resting-state network dynamics that might be “like a ‘special patient group with a vestibular imbalance’ but without lesions in the inner ear or central nervous system” (Boegle et al 2016, p. 420).

However, do these functional modulations by MVS also alter behavioral responses? Would the mere exposure to a 3T MRI scanner already produce the characteristic biases of spatial orientation and attention that we know to occur with vestibular stimulation? For example, in healthy subjects, caloric vestibular stimulation (CVS) of one external ear canal can provoke “neglect-like” behavioral phenomena. Stroke patients with spatial neglect no longer explore or orient towards large parts of space contralateral to their lesion and therefore neglect contralesionally located people or objects (Heilman et al 1983, Karnath 2015, for a review). In healthy volunteers, CVS not only induces a VOR, but also a tonic shift of the average horizontal eye position (Abderhalden 1926, Jung 1953). A further consequence of CVS in healthy subjects is a tonic bias of head orientation around the yaw axis (Karnath et al 2003). Both of these orientation biases resemble those observed in neurological patients suffering from spatial neglect (cf. Fruhmann-Berger & Karnath 2005): already at rest, i.e. when doing nothing, these patients’ head and eyes are tonically deviated towards the ipsilesional side. Moreover, CVS in darkness even induces a bias in subjective straight-ahead (SSA) in healthy subjects, mimicking the bias of the SSA in neglect patients towards their ipsilesional side (e.g. Karnath et al 1994, Chokron & Imbert 1995, Kapoor et al 2001). Finally, CVS biases overt spatial attention. This is obvious from the exploratory scan path in healthy subjects during visual search (Karnath et al 1996). Also this CVS-induced horizontal bias in spatial attention is resembled by the spontaneous, asymmetrical, spatially biased exploratory behavior of neglect patients (Karnath et al 1996, Karnath et al 1998), leading to “spatial neglect” of one side of the surrounding scene. Conversely, it is possible to compensate the spontaneous biases of neglect patients through CVS (Rubens 1985, Karnath 1994b, Vallar et al 1995, Karnath et al 1996, Rode et al 1998; for review e.g. see Rossetti & Rode 2002).

In the present study, we asked whether these behavioral effects on spatial orientation, exploration and attention known from CVS are likewise induced by MVS through exposure to a static magnetic field of a 3T MRI scanner. We hypothesized that MVS should induce a horizontal bias in both the exploratory scan path during visual search and in the SSA of healthy subjects, resembling the horizontal biases of neglect patients.

## Results

To address our research question we analyzed oculomotor behavior of 17 healthy subjects inside and outside a standard 3T MRI scanner (Siemens MAGNETOM Prisma). All subjects provided their informed consent according to the guidelines of our institutional ethics board.

Parts of our procedures resembled that of typical fMRI experiments (compare supplemental Figure S1 and Methods for further details): We placed our subjects with their back on the scanner table and their head rested inside a 20-channel head-coil. A mirror was mounted on top of the head-coil to provide subjects with an indirect view on our custom-made black “visual search-screen”. The screen was placed behind the coil inside the scanner bore at about 110 cm viewing distance. Various glass-fiber-cables penetrated the screen at discrete locations and allowed us presenting small visual stimuli by feeding these cables with LED light signals. Maximal target distance from the screen center amounted to ±12° visual angle in the horizontal direction and to −5° to 6° in the vertical direction. An IR-video-camera unit was mounted next to the mirror to record a video-signal of subjects’ right eye. The study was conducted in complete darkness by extinguishing all light sources and covering subjects with a black blanket. This was important, as any visible stimulus that would be present in subjects’ visual field, could help them to suppress a MVS-induced VOR as well as to explore the visual scene and to perform the SSA task in an unbiased fashion.

To probe for the putative MVS-effects on behavior and attention we designed an experiment comprising of three consecutive phases: (i) an initial *“outside 1”* phase with our subjects on the scanner-table at its maximal horizontal displacement from the scanner center (head coil at ~ 125 cm horizontal [0 cm vertical] displacement); the strength of the magnetic field in this position is roughly 10 times smaller than at the scanner center (compare supplemental Figure S1); (ii) a subsequent *“inside”* phase with the subject’s head being in the center of the scanner bore where the strength of the magnetic field is 3 Tesla; (iii) as well as a final *“outside 2”* phase that was identical to *outside 1*. Accordingly, any MVS effect on behavior or attention that we would observe during these three phases should be stronger for *inside* as compared to the two *outside* phases. Each phase started with an initial *“calibration task”* for eye-tracking (see Methods section for details). We next instructed subjects to fixate at a dim light-point presented in the center of their visual field for 5 seconds. Then the dim light-point was switched off and the subjects had to “look straight ahead” for 60 seconds in complete darkness (*“looking straight-ahead task”*). Finally, we asked subjects to find and fixate transient light stimuli. During this *“visual search task”* we presented 6 search targets (each slowly fading-in over a period of 5 seconds). Locations of search targets were pseudorandomized across phases and between participants. The presentation of these stimuli only served to maintain the subjects’ motivation to search for possible targets. Our interest was the spatial distribution of subjects’ overt spatial attention, as assessed by their exploratory scan path in the absence of any visual targets. Thus, for most of the time (i.e. 140seconds; total duration of visual search task: 172 seconds), no visible target was present and subjects were searching in complete darkness. For data analysis we discarded the time periods with targets present (plus an additional grace period of 5 seconds after a light stimulus was turned off; this was done to prevent carry-over effects from prior target-fixation). The search task always ended with a central light stimulus presented during the last 2 seconds (also not considered for data analysis). Please refer to supplemental Figure S2 for an overview over all tasks and phases of our experiment.

Figure 1 illustrates the resulting eye-data of an exemplary subject throughout all tasks and phases. It shows that during the *inside* phase the typical saw-tooth-like eye movement pattern, which is characteristic for a vestibular nystagmus, was observed. It consisted of a slow rightward VOR that was accompanied by fast resetting saccades in the opposite direction. This prototypical nystagmus-pattern was largely reduced for the *outside 1* phase and it was practically absent for the *outside 2* phase (also compare our supplemental video). To express the strength of the VOR quantitatively, we calculated the median de-saccaded horizontal eye-velocity (starting 5 seconds after the offset of the central light stimulus). This estimate is reflected by the blue lines in the respective horizontal eye-velocity time-courses shown in the middle column of Figure 1.

**Figure 1.**
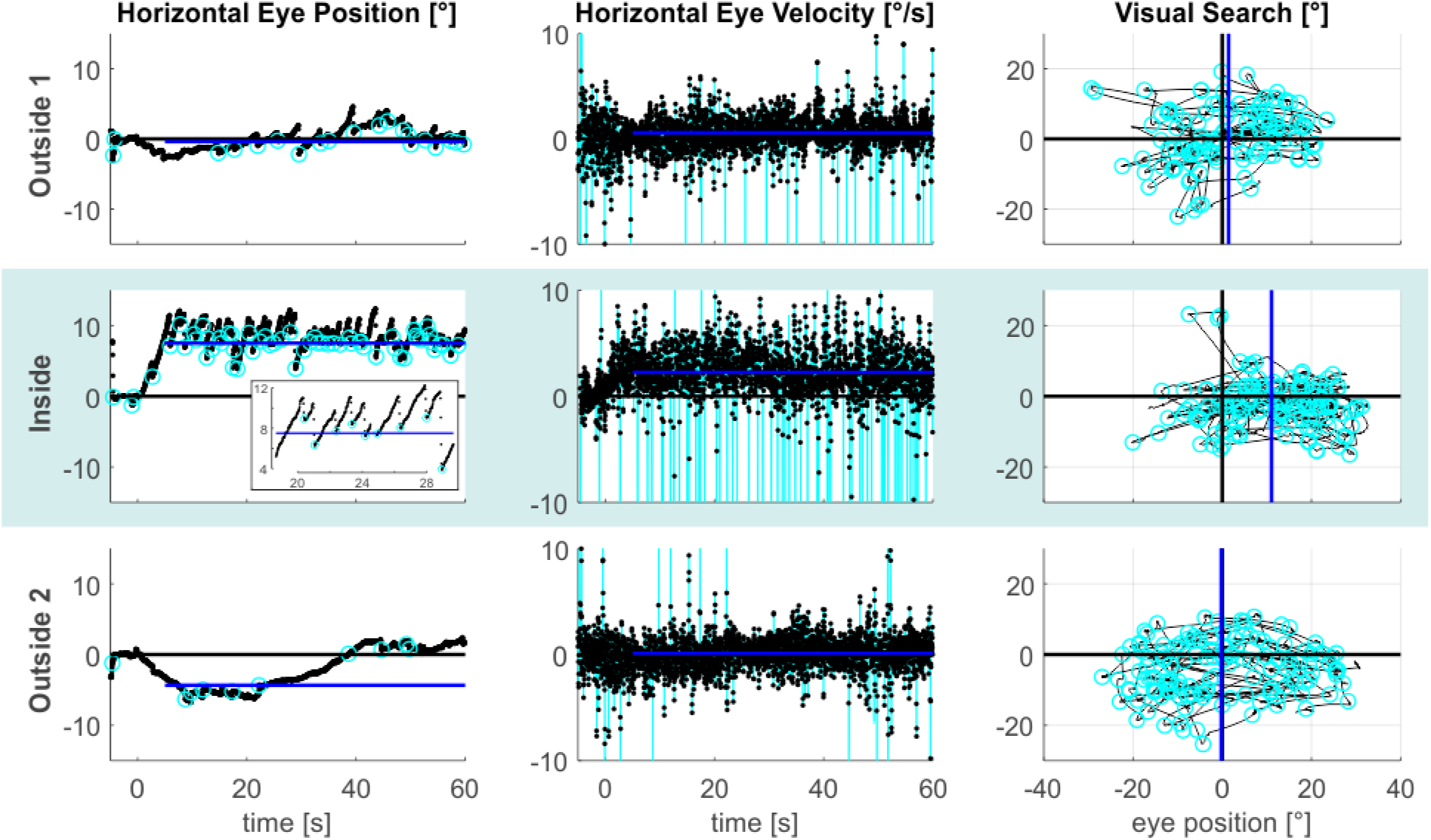
MVS-induced oculomotor behavior. The figure provides exemplary data from an individual subject. The three task phases are represented by individual rows. Horizontal eye position data from the looking straight-ahead task are shown in the left column. The figure inset additionally expands parts of the *inside* time course to better illustrate the alternation between the slow phase VOR towards the right and the fast resetting saccades directed in the opposite direction. Saccade endpoints are depicted by the cyan circles and the blue lines reflect the respective SSA estimates. Corresponding horizontal eye velocity traces of the looking straight ahead-task are shown in the middle column. Note that the cyan peaks in these time-courses refer to individual saccades, which have been removed from the eye velocity records to allow estimation of slow-phase velocity in isolation (the blue lines indicate the respective horizontal VOR-estimates). Finally, 2D eye-position eye data from the visual search task are depicted in the rightmost column (time periods with search targets plus 5 seconds are excluded). The horizontal center of visual search is depicted by the blue vertical lines, reflecting the median of horizontal saccade endpoints (cyan circles). Positive values indicate the rightward/upward direction.

The same qualitative effects were present in all of our 17 subjects, as is shown in Figure 2. The average slow eye velocity for the *outside 1* phase was 0.30°/s on average (±0.49°/s standard deviation [SD]) and increased to 1.42°/s (±1.49°/s SD) for the *inside* phase. After removing the subjects from the scanner bore to *outside 2*, average slow eye velocity decreased to −0.01°/s (±0.40°/s SD). The within-subject differences in velocity between *outside 1* and *inside* and between *inside* and *outside 2* were significant (one-tailed paired t-test: t(16)=6.62, p<0.001, g_1_[CI_95%_]=1.60 [0.87, 2.32] and t(16)=6.69, p<0.001, g_1_[CI_95%_]=1.62 [0.88, 2.35], respectively). Average horizontal eye velocity during *outside 1* was also slightly larger as compared to *outside 2* (two-tailed paired t-test: t(16)=3.08, p=0.007, g_1_[CI_95%_]=0.75 [0.20, 1.28]). As expected (Otero-Millan et al 2017), the MVS effects affected only horizontal eye movements and were absent in the vertical direction (outside 1: −0.29°/s±0.56°/s SD; inside: −0.29°/s±0.60°/s SD; outside 2: −0.21°/s±0.58°/s SD; statistical comparisons with two-tailed paired t-tests yielded no significant differences in vertical eye velocity between any of the three task phases [p>0.20]). In summary, these data agree with earlier observations of an MVS-induced horizontal VOR (for review see (Ward et al 2019) and provide additional evidence for the consistent presence of this effect while being inside a 3T MRI scanner (cf. Boegle et al 2016).

**Figure 2.**
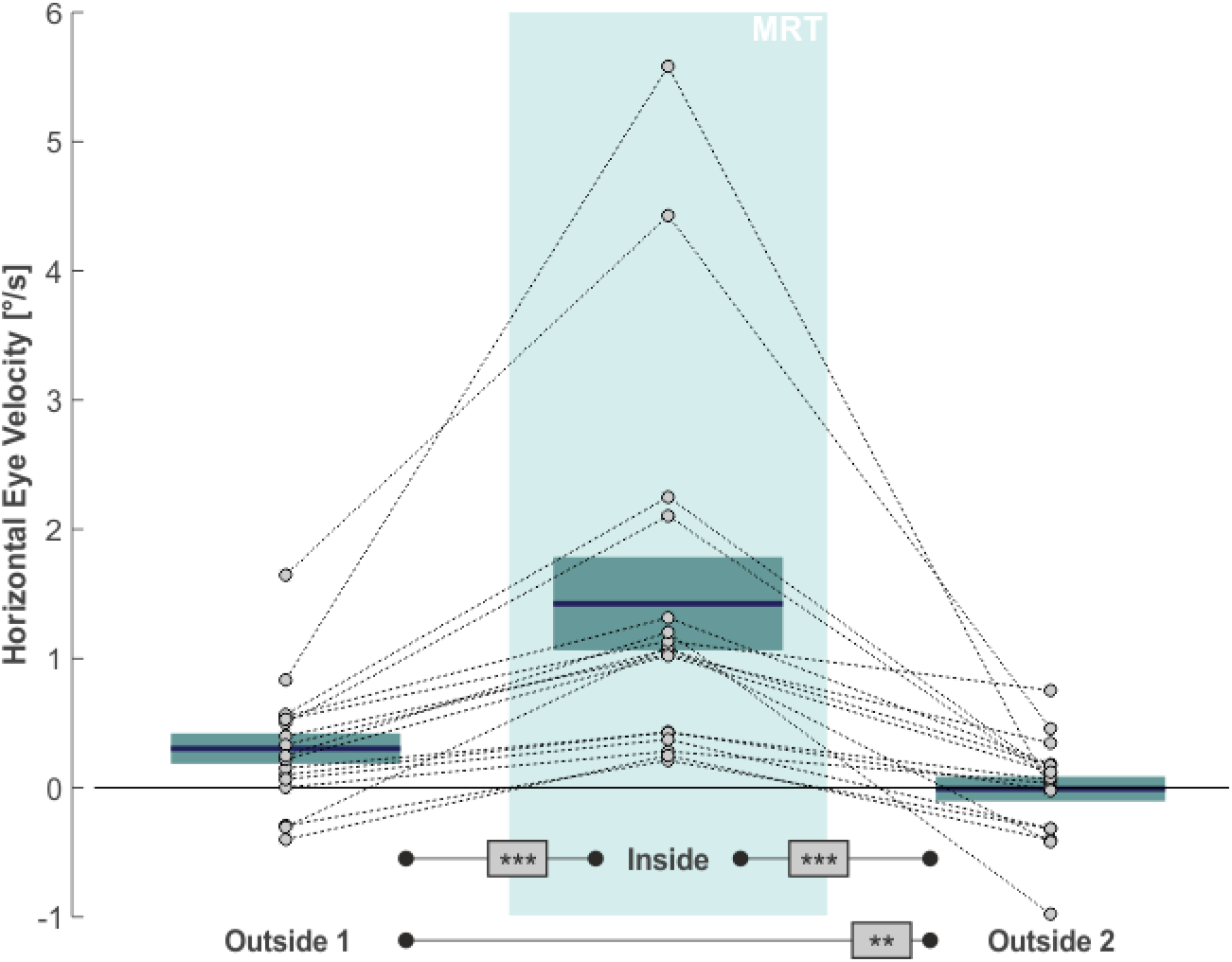
MVS-induced changes in de-saccaded horizontal eye velocity. Horizontal eye velocity of individual subjects (circles, N=17) are shown for all three task phases. Positive values denote rightward motion. The blue lines and the boxes reflect across subject means and standard error (SE; to account for our within-subject design, we removed the between-subject variance for calculation of the SE according to the procedure described by Masson & Loftus (2003)). Results of pairwise statistical comparisons are indicated: ** 0.001≤p<0.01; *** p<0.001.

We next asked whether MVS also affects the distribution of spatial attention in our visual search task. The rightmost column in Figure 1 demonstrates a 2D-plot of our exemplary subject’s eye position during search, including only those time epochs when no target stimulus was presented (cf. above). While the area of visual search for both *outside* phases was centered on the middle of the screen (which was aligned to the subject’s body midline), the search space clearly deviated to the right when the subject was *inside*, while parts on the far left of the search-space were now neglected. This horizontal bias of exploratory behavior was not a mere consequence of the rightward VOR inside the scanner but was likewise reflected by the distribution of the endpoints of the saccades guiding visual search: the mean horizontal position of these saccade endpoints (indicated by the blue vertical lines in the rightmost plots in Figure 1) almost perfectly resembled the center of the search screen for the *outside 1 & 2* phases, while it was clearly shifted to the right for the *inside* phase. Across our 17 subjects, statistical analysis of subjects’ exploratory spatial behavior (Fig. 3) revealed that the average horizontal mean of visual search was at 1.65° (±4.40° SD) and at 1.01° (±3.98° SD) for the *outside 1 & 2* phases, respectively, while *inside* this measure was significantly shifted to 4.69° (±4.61°) towards the right (one-tailed paired t-tests; *outside 1* vs. *inside*: t(16)=−5.12, p<0.001, g_1_[CI_95%_]=1.24 [0.59, 1.87]; *inside* vs. *outside 2*: t(16)=4.38, p<0.001, g_1_[CI_95%_]=1.06 [0.45, 1.65]). The measures did not significantly differ between phases *outside 1 & 2* (two-tailed paired t-test: t(16)=0.76, p=0.46, g_1_[CI_95%_]=−0.18 [−0.66, 0.30]).

**Figure 3.**
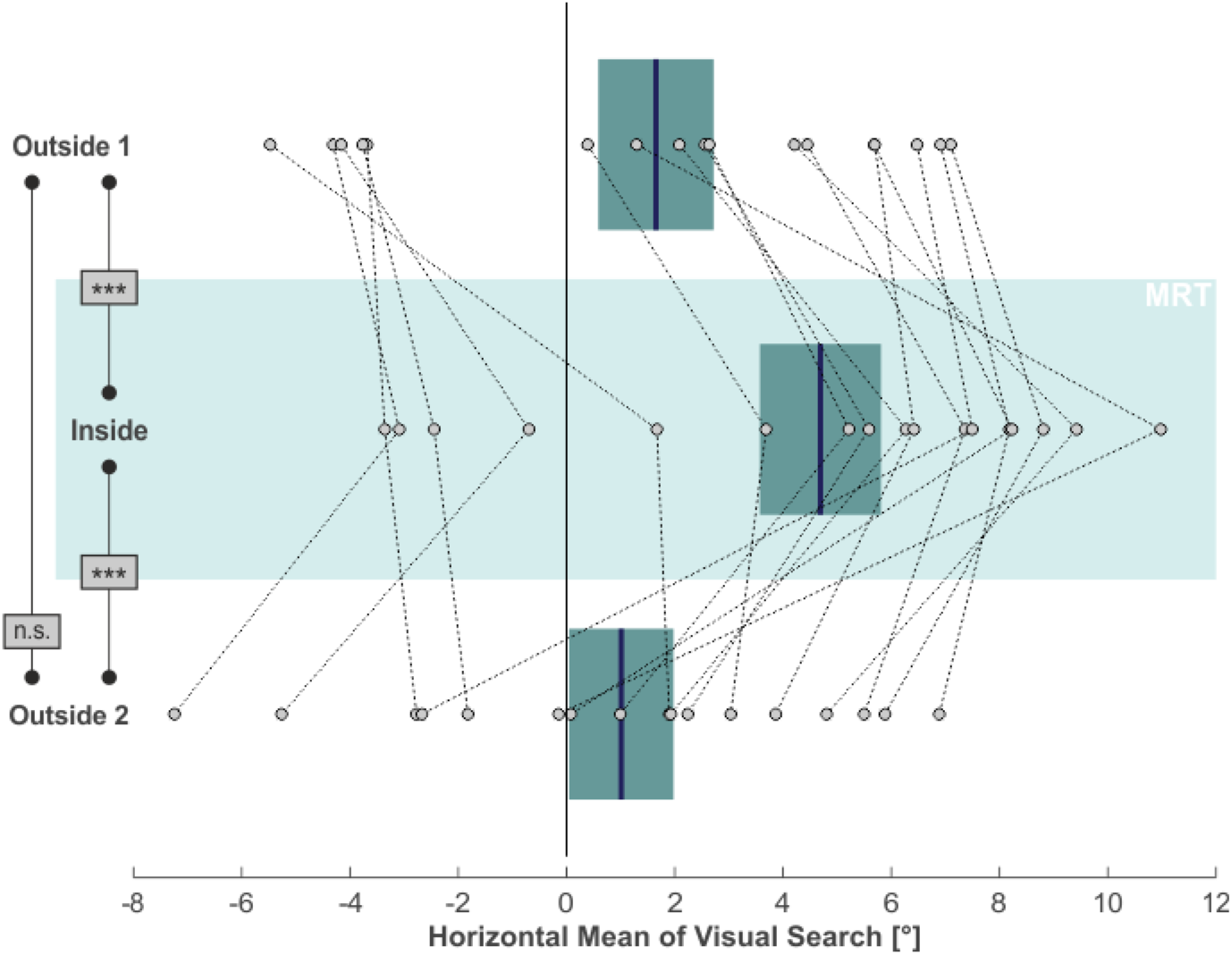
MVS-induced changes in visual search. Individual subjects’ horizontal mean saccade endpoints during visual search (in the absence of targets) are depicted as individual circles and for all three task phases (N=17; positive values = right). The blue lines and the boxes reflect across subject means and SE (to account for our within-subject design, we removed the between-subject variance for calculation of the SE according to the procedure described by Masson & Loftus (2003)). Results of pairwise statistical comparisons are indicated: n.s. not significant (p≥0.05); *** p<0.001.

Finally, we asked whether the MVS-induced spatial bias would also affect the looking straight ahead task. To this end we again focused on the horizontal distribution of saccade endpoints during this task, as they reflected subjects’ voluntary attempt to continue looking straight ahead. For each subject and for each task phase we calculated the median horizontal position of saccade endpoints (starting 5 seconds after the offset of the central light stimulus) as a proxy for the subjective straight-ahead (SSA). This measure is illustrated by the blue lines in the leftward column of Figure 1 for our exemplary subject. Similar to the horizontal shift of exploratory search behavior across task phases, also the looking straight ahead measure shifted. Initially, the SSA was close to the midline during the *outside 1* phase. It shifted to the right for the *inside* phase and then back towards the left (with some overshoot) for the *outside 2* phase. Across our 17 subjects, the SSA likewise exhibited a significant rightward shift from the *outside 1* (1.79°±5.39° SD) to the *inside* (4.76°±4.02° SD) phase (one-tailed paired t-test: t(16)=−2.38, p=0.015, g_1_[CI_95%_]=0.58 [0.05, 1.08]; compare Fig. 4). This effect vanished after removing subjects from the scanner in the *outside 2* phase (0.26°±4.37° SD; one-tailed paired t-test: t(16)=3.56, p=0.001, g_1_[CI_95%_]=0.86 [0.29, 1.42]). There was no significant difference between the SSA for phases *outside 1 & 2* (two-tailed paired t-test: t(16)=1.17, p=0.26, g_1_[CI_95%_]=−0.28 [−0.76, 0.21]).

**Figure 4.**
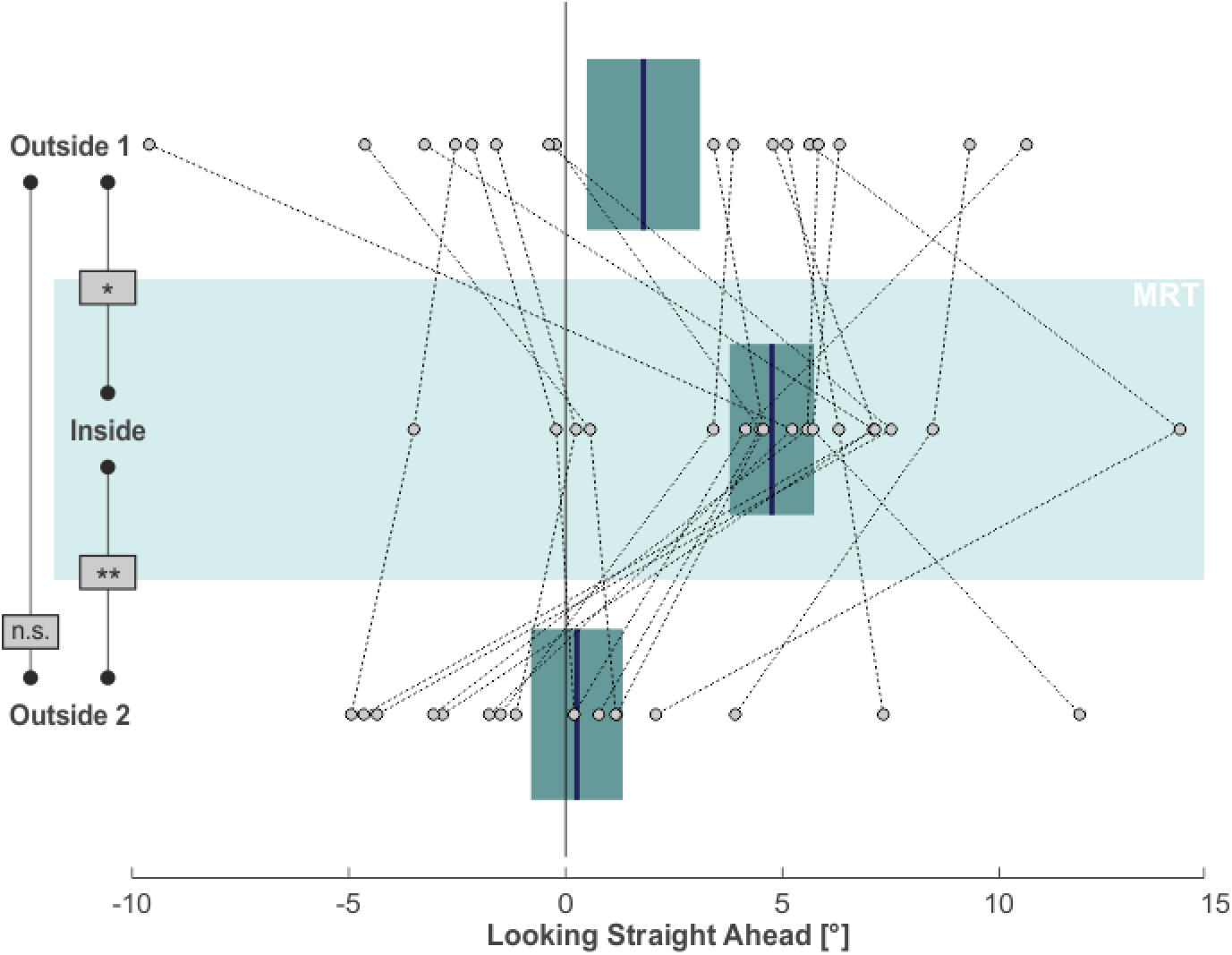
MVS-induced changes in subjective straight ahead. Individual subjects’ SSA, namely the median horizontal saccade endpoint when trying to look straight ahead, is depicted by individual circles for all three task phases (N=17; positive values = right). The blue lines and the boxes reflect across subject means and SE (to account for our within-subject design, we removed the between-subject variance for calculation of the SE according to the procedure described by Masson & Loftus (2003). Results of pairwise statistical comparisons are indicated: n.s. not significant (p≥0.05); * 0.01≤p<0.05; ** 0.001≤p<0.01.

## Discussion

Like previous studies (for review see Ward et al 2019), we observed a MVS-induced tonic direction-specific (rightward) horizontal VOR when lying inside a 3T MRI scanner. As compared to the *outside 1&2* phases, this effect was present in every single subject. The velocity of the VOR inside the MRT amounted to 1.42°/s during our looking straight-ahead task. Our task instruction to fixate an “imaginary target”, namely into the direction of a previously illuminated target LED in the medial plane, probably reduced VOR-velocity by about 25% as compared to a situation without such imaginary fixation (cf. Schmäl et al 2000). Given this latter consideration, our estimate well corresponds to the roughly 2°/s VOR reported in an earlier MVS study, which used a comparable experimental setting at 3T but without imaginary fixation (Boegle et al 2016) (for further discussion of our VOR results, please refer to our supplemental discussion).

Our experiment further revealed that lying inside the 3T MRI scanner also led to a spatially biased neglect-like behavior when subjects were trying to look straight ahead and when they were performing visual search. First of all, we would like to point out that the respective biases in the looking straight ahead task and the visual search task were not a mere reflection of the tonic VOR-induced drift of subjects’ eyes towards the right. The critical behavioral measure in both tasks refers to the distribution of endpoints of subjects’ voluntary saccades while they were trying to follow task instructions. The passive drift of subjects’ eyes due to the MVS-induced VOR does not influence this measure of active spatial behavior (for further discussion also cf. Karnath et al 1996, p.340). Hence, the reported shifts inside the scanner reflect a true spatial bias in overt spatial attention and active goal-directed behavior: (i) in the visual search task the center of subjects’ visual scan-path (in the absence of any visible target) was shifted by roughly 3.4° to the right; (ii) in the looking straight ahead task, the average endpoint of subjects’ saccades in darkness likewise deviated towards the right, namely by about 3.7°.

Our MVS results resemble the effects of caloric vestibular stimulation on spatial attention and orientation: After CVS, healthy subjects’ estimates about SSA, quantified by orienting a laser-spot into this direction, shifted horizontally (e.g. Karnath et al 1994, Karnath et al 1996). Moreover, CVS likewise leads to a horizontal shift of healthy subjects’ scan path during visual search (Karnath et al 1996). These CVS-induced deviations ranged roughly between 4° and 9° and thus were slightly larger than the MVS-induced spatial bias observed in the present experiment. Importantly, the CVS-induced biases in spatial behavior also surface when trying to manually point straight ahead (Schmäl et al 2000) and during manual spatial exploration (Karnath et al 2003), respectively. This suggests that CVS − and probably also MVS − does induce changes in spatial representations for goal-directed actions that are effector-independent. These CVS/MVS-induced alterations of spatial behavior are most likely of cerebro-cortical origin. Notably, there exists no “primary vestibular cortex” in the strict sense but rather an interconnected multimodal “cortico-vestibular system” with the parieto-insular vestibular cortex (PIVC) at its core (e.g. see Guldin & Grüsser 1998, Karnath & Dieterich 2006, Lopez & Blanke 2011 for review). The areas that make up this system further entail neighboring parts in the temporo-parietal junction, posterior parietal cortex, superior temporal cortex, as well as regions in the anterior insula, retroinsular regions, somatosensory cortex, cingulate cortex, premotor cortex, and in the hippocampus (Guldin & Grüsser 1998, Kahane et al 2003, Karnath & Dieterich 2006, Lopez et al 2012, zu Eulenburg et al 2012, Mazzola et al 2014, Frank & Greenlee 2018). Neurons in part of these areas have been shown to integrate vestibular cues (along with neck proprioception and information about eye position) in a way that allows them to represent visual action goals in reference to the body (e.g. Andersen et al 1999, Cohen & Andersen 2002, Chen et al 2018). This body-centered, egocentric representation of spatial information underlies, e.g., attentional orienting and the guidance of our goal-directed behavior including saccades, reaching, or heading etc. (see Karnath 2001 for review). Such “body referencing” has also been reported in human functional imaging studies (e.g. Bottini et al 2001, Galati et al 2010, Schindler & Bartels 2013, Saj et al 2020). It thus is conceivable that tonic MVS (as well as CVS, etc.) introduces systematic biases in these spatial action-representations, leading to the reported alterations of subjective straight ahead and spatial exploration (also compare Laurens & Angelaki 2018). Similarly, altered body referencing was suggested to explain the horizontal shifts in spatial behavior present in spatial neglect (Karnath 1994a; see Karnath 2015 for a review). Moreover, the pattern of lesions exhibited by patients with spatial neglect is largely overlapping with the multimodal cortico-vestibular system and exhibits a common lateralization (Karnath & Dieterich 2006, Karnath & Rorden 2012, Dieterich & Brandt 2015). Hence, MVS induces neglect-like alterations of spatial behavior in healthy subjects and these alterations are supposedly due to its interference with the cortico-vestibular system, the same system that is typically affected in patients suffering from spatial neglect.

Given the aforesaid, the question arises whether MVS might be a viable option for treatment of spatial neglect. In this context, the observations that healthy subjects exposed to the static magnetic field of a MR scanner developed a persistent horizontal nystagmus that slowly diminished but did not extinguish (Roberts et al 2011, Jareonsettasin et al 2016), is highly interesting since the effect of, e.g., CVS only lasts for several minutes. Thus, MVS is a potential noninvasive and comfortable way to continuously stimulate the labyrinth, which may be a promising new treatment option for neurological patients with spatial neglect. We expect that neglect patients would benefit from MVS to an extent that is at least comparable to the effects which have been reported for CVS (cf. above) and other types of vestibular stimulation (see Kerkhoff & Schenk 2012 for a review). The tonic nature of MVS might help to induce longer-term plastic changes in patients’ pathologically altered spatial representations (cf. Jareonsettasin et al 2016).

Finally, as was already pointed out by Roberts et al. (Roberts et al 2011), we want to emphasize that MVS effects – like the ones presented in our study – are likely present during any fMRI study at least at 3T. Boegle and coworkers (2016) demonstrated that even for the default positioning of subjects inside a 3T MRI scanner an MVS-induced VOR was present in most subjects; moreover, they showed that MVS had an impact on resting state activity in the default mode network and in particular in those areas related to vestibular and oculomotor function. In their follow-up work these authors further demonstrated a MVS-induced modulation of visual-vestibular network activity (Boegle et al 2017). Our study suggests that MVS-effects on spatial attention, perception and behavior during task-related fMRI must be considered in addition.

## Methods

### Subjects

We recruited seventeen healthy subjects for our experiment (5 males; average age 25.1±5.0 years SD). All subjects were right-handed, had normal or corrected to normal vision, and provided their informed consent according to our institutional ethics board guidelines prior to our experiment. Sample size was guided by two separate power analyses (α=0.025; power=97.5%, each) informed by the studies by Boegle et al (2016; 3T MVS-induced VOR; data from their figure 1a) and Karnath et al (1996; changes of SSA due to CVS and neck muscle vibration; data from their table 1), respectively. Both analyses suggest a minimum sample size of 13 subjects (one-tailed one-sample/paired t-test). To account for putative subject dropouts, we decided to recruit four additional subjects, amounting to a total sample size of 17 subjects.

### MRI Setting

We used a 3T Siemens MAGNETOM Prisma MRI Scanner to apply MVS. No radio frequency (RF) or gradient coil fields were applied. Given standard subject positioning, the magnetic field vector of our MRI system pointed from subjects’ toes to their head. Note that this is the exact opposite direction as compared to the situation in the study of Roberts and colleagues (2011). Accordingly, our MVS-induced VOR effects were in the opposite horizontal direction as compared to theirs. A map of the magnetic fieldlines of our system is provided in Supplemental Figure S1. This figure also details the location of our subjects’ head for the *outside 1&2* phases as well as for the *inside* phase.

We tried to maximize the effects of MVS on the horizontal canals by positioning subjects in a way that their head would tilt slightly backwards inside the head-coil (Siemens Head/Neck 20 A 3T Tim Coil). According to the results reported by Roberts et al. (2011) and Boegle et al. (2016), such “pitching-up” of subject’s head can increase MVS-effects on the VOR. Based on these authors’ results we approximated a head pitch angle of about −30° (canthus-tragus-line vs. vertical) to consistently achieve strong MVS-effects in all subjects. To this end we placed various cushions underneath subjects’ neck. We also placed cushions to the left/right of subjects’ head inside the head-coil to prevent any head movements.

### Stimulus Generation

An overview of our general experimental design is depicted in Supplemental Figure S2. A WIN10 laptop PC was used to control our experiment through custom scripts using MATLAB R2015b 32bit (MathWorks) in combination with Cogent 2000 and Cogent Graphics (by FIL, ICN, LON at the Wellcome Department of Imaging Neuroscience, University College London) as well as with the MATLAB Support Package for Arduino. Light stimuli were generated through 8 red LEDs (Type: L-513HD; Dropping resistor: 58kΩ) using an Arduino UNO R3-compatible microcontroller (Funduino UNO R3) that was controlled by MATLAB. The intensity of six of those LEDs could be adjusted using the Arduino’s pulse code modulated (PCM) analog voltage output (0V-5V). The remainder of the LEDs could only be switched on/off (0V or 5V). The LED-light was fed into a bundle of 8 glass-fiber cables that went from the MRI-control room into the scanner room at predefined locations within the subject’s search field (fiber-diameter: 1mm ≙ 0.05° visual angle; viewing distance = 1.1 m; note that viewing distance corresponded to the “resting state” of eye convergence in darkness [Owens & Liebowitz 1980]). By this means we were able to deliver dim light stimuli through the fiber-ends inside the scanner. Apart from these transient light stimuli, subjects remained in complete darkness.

During the visual search task the voltage of the 6 search target LEDs was PCM-modulated across the 5 seconds presentation time for a given target, starting with 0.1V while doubling voltage in 1 second steps up to 1.6V. Through this manipulation we slowly increased the visibility of the search targets to increase the likelihood that subjects would find them. Remember that for most of the time (i.e. 140 seconds) no target was visible at all. The final central target at the end of the visual search task (as well as all calibration targets and the initial central target of the looking straight ahead task) were always presented with full intensity (5V). The six search targets were at the following x/y–locations (values denote visual angle; positive values represent rightward/upward directions, respectively): −6°/5°; 6°/5°; −6°/−5°; 6°/−5°; −12°/−1°; 12°/1°. Importantly, in our visual search task the order of targets was pseudorandomized across task phases and between participants. The final central search target was always at 0°/0°. The same central target was shown during the initial 5 seconds of the looking straight ahead task. Finally, the following 5 targets served for eye-calibration: 0°,0°; −6°/5°; 6°/5°; −6°/−5°; 6°/−5°.

### Manual Responses

During the search task, subjects were asked to find and fixate any target that they would spot. In addition, we asked them to press a button on a MRI-compatible response pad (5-Button Diamond Response Pad; Current Designs), whenever they found a search target. This instruction chiefly served to keep subjects’ search motivation high throughout the search period. As mentioned above, we were not interested in target hits but in subjects’ exploratory scan paths when no target was present. Thus, we did not systematically analyze subjects’ responses. Still we provide an estimate of response performance for the *inside* and *outside 2* phases, in which we collected data in the vast majority of subjects (n=14 and n=15 [manual responses were not reliably recorded in all subjects for technical reasons]): hit-rates were 91.7% ± 17.0% SD and 93.3%±13.8% SD, and average reaction times were 3236ms±698ms SD and 2995ms±697ms SD, respectively.

### Eye Tracking

The position of subjects’ right eye was monitored at 50 Hz sampling rate with an MR-compatible camera with integrated infrared LED illumination (MRC Systems; Model: 12M-i IR-LED). We used the ViewPoint Monocular Integrator System and the View Point Software (Arrington Research, software version 2.8.3.437) to digitize the eye-camera video and to obtain uncalibrated eye-position data (by means of dark pupil tracking). Eye-tracking was realized on a different WIN PC that was remote-controlled through the ViewPoint Ethernet-Client running on our laptop PC. Eye movement analyses were performed off-line using custom routines written in MATLAB R2017b (MathWorks). In brief, eye position samples were filtered using a second-order 10 Hz digital low-pass filter. A 5-point calibration was performed based on the data obtained in our calibration task. In addition, we compensated for any tonic eye position offset in all other tasks as well. Compensation was performed separately for each task and phase, namely by removing the (average) difference in position between the visible fixation/search target(s) and eye position during target fixation(s) from the eye position record. Eye velocity was calculated based on 2-point differentiation of our eye position data. Saccades were detected using an absolute eye velocity threshold of >15°/s. Saccade-onset was defined as the sample prior threshold-crossing. Saccade-offset was defined accordingly, namely as the first sample after eye velocity dropped below the threshold (compare corresponding saccade endpoints depicted as cyan circles in Figure 1). Time periods with blink artifacts were excluded from analyses. To obtain our velocity estimates for MVS-induced VOR we used de-saccaded horizontal/vertical eye-velocity traces (time-periods from saccade-onset to saccade-offset were treated as missing values; compare middle column of Figure 1).

### Statistical Analyses

Apart from the de-saccaded eye velocity estimates obtained at the *inside* phase, all other data of interest (and their respective paired differences between phases) were normally distributed (Shapiro-Wilk-Test; p≥0.01). Accordingly, we applied paired t-tests (alpha=5%) to these latter data, namely between *inside* and *outside 1*, between *inside* and *outside 2*, and between *outside 1 & 2* phases, respectively. For statistical comparison of the de-saccaded eye velocity estimates during the *inside* phase with both *outside 1&2* phases, we log-transformed the respective paired differences to ensure normal distribution (Shapiro-Wilk-Test; p≥0.01). These log-transformed data were further analyzed by means of one sample t-tests (alpha=5%). Due to clear directional hypotheses concerning MVS-effects on behavior between *inside* and *outside 1 & 2* phases we applied one-tailed tests. Two-tailed tests were applied when comparing *outside 1 & 2* phases and when analyzing vertical eye velocity, which should be largely unaffected by MVS. Note, that all reported p-values also survived Bonferroni-correction for multiple testing within each measure of interest (adjusted alpha=1.7%). Effect size estimates (Hedges g_1_±CI_95%_) were calculated using the Matlab Toolbox ‘Measures of Effect Size’ (Version 1.6; by H. Hentschke and M.C. Stüttgen).

### Data sharing and code availability

All data and custom code from this study will be made available by the authors upon reasonable request.

## Acknowledgements

This work was supported by the Deutsche Forschungsgemeinschaft (KA 1258/23-1). Daniel Wiesen was supported by the Luxembourg National Research Fund (FNR/11601161). We particularly thank Hannah Rosenzopf and Stefan Smaczny for assisting us during our measurements. We also thank Marc Himmelbach and Michael Erb for their suggestions and help with our setups and for general MRI-support.

## Supplement

**Figure S1.**
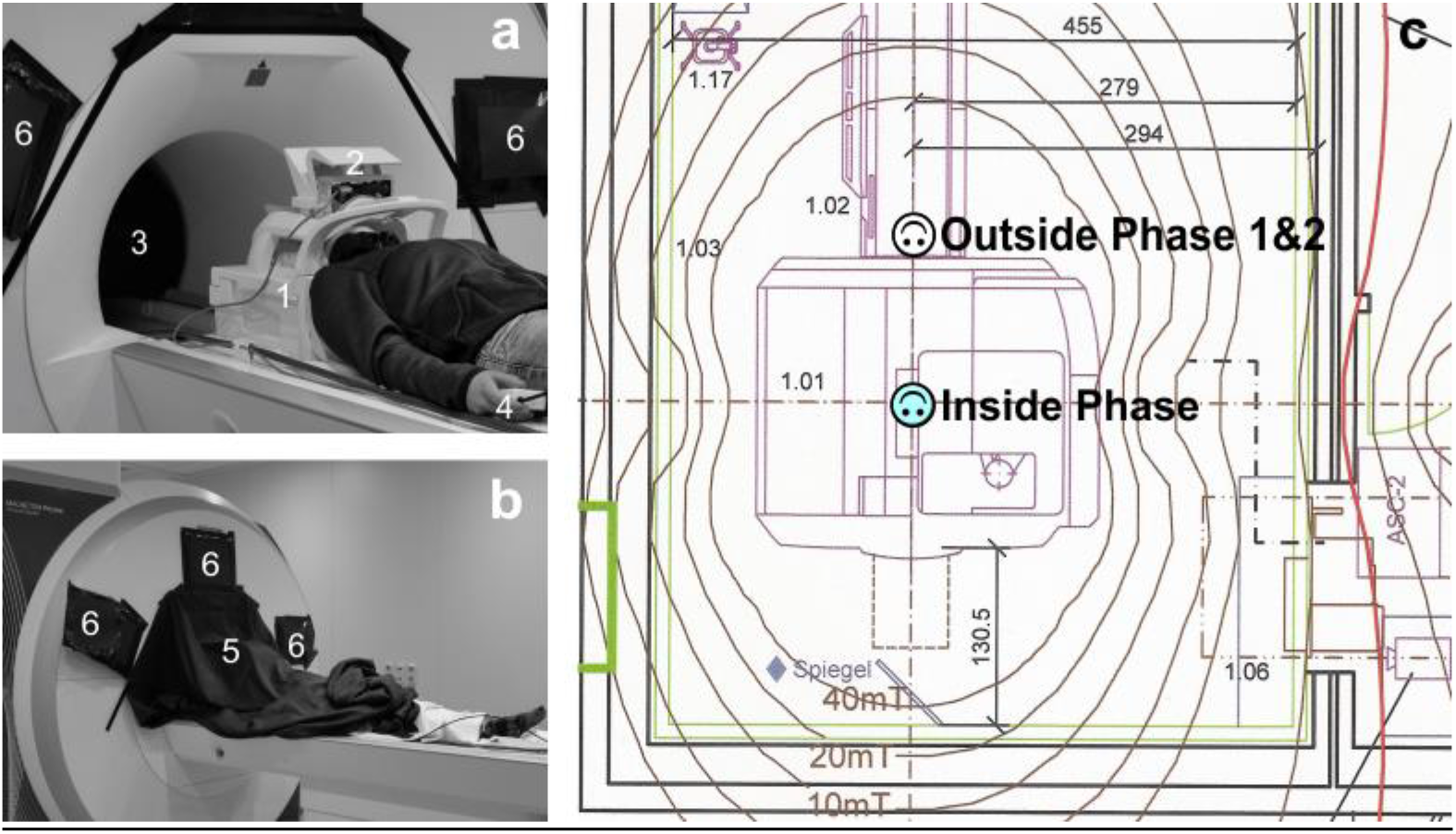
Experimental Setup. The photographs on the left (**a & b**) show the positioning of a subject representative for the *outside 1 & 2* phases. **a** The subject was lying with the head inside the head-coil (1); a mirror and eye-tracking system were mounted on top of the coil (2). The search screen (3) was inside the MRI bore and could be viewed via the mirror. The subject was holding a MRI-compatible response box in the right hand (4). **b** After positioning of the subject, we covered her with a black blanket that was attached to the front of the MRI scanner (5). We further covered any light sources inside the scanner room with black cardboard (6). Note that we also covered the opening of the scanner bore in the back (not shown) and we also turned off any lights inside the scanner room and in all neighboring rooms. These measures allowed us to guarantee a completely dark environment for our experiments. **c** The illustration on the right shows a field-line drawing from our scanner manufacturer and depicts the position and orientation of subject’s head for the respective experimental phases.

**Figure S2.**
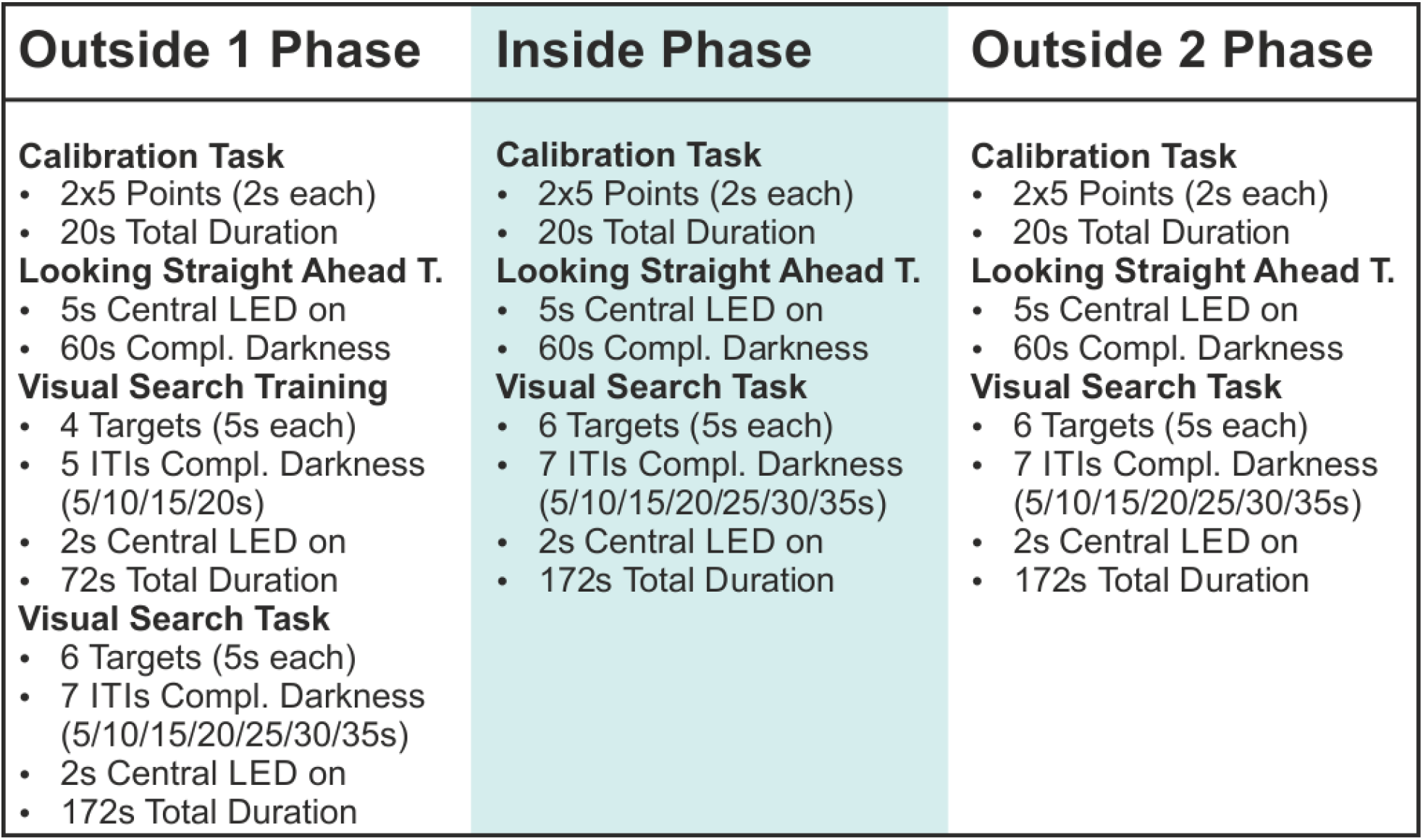
Experimental Design. All tasks are listed in their respective order and all relevant temporal parameters are provided. Average duration for the completion of all tasks (including additional breaks and instructions) amounted to 505s±58s SD in the *outside 1* phase. Due to the additional search training in this phase, this duration was larger than for the *inside* phase and for the *outside 2* phase, which lasted 349s±15s SD and 347s±12s SD, respectively. The average inter-phase-intervals amounted to 233s±87s SD between *outside 1* and *inside* phases as well as 241s±67s SD between the *inside* phase and *outside 2* phase. The time of subject’s head being in the center of the MRI-scanner was about 10 minutes on average.

## Supplemental Movie

**Supplemental Movie.** A supplemental movie can be assessed online: https://my.hidrive.com/share/ijtl.djm9b. This supplemental movie was taken after completion of our experiment. It serves to illustrate the MVS-induced effects on eye movements in complete darkness. There was no obvious horizontal nystagmus when the subject was outside the scanner (o*utside 1*). A pronounced horizontal nystagmus was present *inside* the scanner (and already when the subject was being moved into the scanner). The horizontal nystagmus was alleviated when moving the subject outside the scanner. At the *outside 2* phase virtually no nystagmus was present.

## Supplemental Discussion

Different to the phasic acceleration-patterns that characterize head motion in most natural settings, MVS occurs tonically – i.e. as if the rotation of the head would be accelerating continually. In fact, a MVS-induced VOR well compares to that induced by putting a subject on a rotating chair that is constantly accelerating (compare Jareonsettasin et al 2016). It is perhaps not surprising (also due to conflicting afferent inputs signaling spatial stability, such as from the otoliths etc.) that the nervous system counteracts the MVS-induced tonic imbalance: This is obvious from a slow but incomplete reduction of the VOR over the course of minutes to hours (Jareonsettasin et al 2016). Such “set-point adaptation” is likely to account for the significant decrease in de-saccaded horizontal eye velocity in our *outside 2* phase as compared to *outside 1* (due to the intervening ~10min exposure to the 3T static magnetic field).

## Notes

**The authors declare no competing interest**

https://my.hidrive.com/share/ijtl.djm9b

### Competing Interest Statement

The authors have declared no competing interest.

